# A SARS-CoV-2 mini-genome assay based on negative-sense RNA to study replication inhibitors and emerging mutations

**DOI:** 10.1101/2021.06.28.450211

**Authors:** Thomas Vial, Michael S. Oade, Colin A. Russell, Dirk Eggink, Aartjan J.W. te Velthuis

## Abstract

Severe Acute Respiratory Coronavirus 2 (SARS-CoV-2) is a positive-sense single-stranded RNA virus and the causative agent of the Coronavirus disease 2019 (COVID-19) pandemic. Efforts to identify inhibitors of SARS-CoV-2 replication enzymes and better understand the mechanisms underlying viral RNA synthesis have largely relied on biosafety level 3 (BSL3) laboratories, limiting throughput and accessibility. Recently, replicon systems have been proposed that involve ^~^30 kb RNA-based replicons or large plasmids that express the viral structural and non-structural proteins (nsp) in addition to a positive-sense reporter RNA. Unfortunately, these assays are not user-friendly due to plasmid instability or a poor signal to background ratio. We here present a simple mini-genome assay consisting of a ^~^2.5 kb-long negative-sense, nanoluciferase-encoding sub-genomic reporter RNA that is expressed from a plasmid, and amplified and transcribed by the SARS-CoV-2 RNA polymerase core proteins nsp7, nsp8 and nsp12. We show that expression of nsp7, 8 and 12 is sufficient to obtain robust positive- and negative-sense RNA synthesis in cell culture, that addition of other nsps modulates expression levels, and that replication of the reporter RNA can be inhibited by active site mutations in nsp12 or the SARS-CoV-2 replication inhibitor remdesivir. The mini-genome assay provides a signal that is 170-fold above background on average, providing excellent sensitivity for high-throughput screens, while the use of small plasmids facilitates site-directed mutagenesis for fundamental analyses of SARS-CoV-2 RNA synthesis.

**Importance statement:** The impact of the COVID-19 pandemic has made it essential to better understand the basic biology of SARS-CoV-2, and to search for compounds that can block the activity of key SARS-CoV-2 replication enzymes. However, studies with live SARS-CoV-2 require biosafety level 3 facilities, while existing replicon systems depend on long positive-sense subgenomes that are often difficult to manipulate or produce a high background signal, limiting drug-screens and a rapid analysis of emerging SARS-CoV-2 mutations during the COVID-19 pandemic. To make it easier to study emerging SARS-CoV-2 mutants and screen for inhibitors, we developed a simple mini-replicon that produces a minimal background signal, that can be used in any tissue culture lab, and that only requires four small plasmids to work.

## Introduction

The emergence of severe acute respiratory syndrome coronavirus 2 (SARS-CoV-2), the causative agent of coronavirus disease 2019 (COVID-19), took the world by surprise (1, 2). Diagnostic tests are now used in most countries, and multiple vaccines are available to control the epidemic. In addition, various candidates for intravenous antiviral treatment have been identified or been given emergency approval (3). However, SARS-CoV-2 continues to mutate rapidly, including the rise of so-called Variants-of-Concern (VOC) and cause severe disease in countries where vaccines and intravenous antiviral treatments are not readily available (4, 5). In addition, immune compromised patients need antiviral treatments to recover from SARS-CoV-2 infections. There is therefore still an urgent need for the development of alternative antiviral treatments and a better understanding of SARS-CoV-2 replication. Such efforts may also aid the development and understanding of other respiratory RNA viruses, such as influenza A virus, respiratory syncytial virus, and other coronaviruses.

The SARS-CoV-2 genome consists of 29,865 nt, excluding polyA-tail, of single-stranded positive-sense RNA (1, 2, 6). The virus relies on the activity of the viral RNA polymerase for the replication and transcription of this genome. Similar to SARS-CoV (7, 8), the RNA polymerase activity of SARS-CoV-2 resides in the RNA-dependent RNA polymerase (RdRp) domain of non-structural protein 12 (nsp12) (9, 10). SARS-Cov-2 nsp12 requires nsp7 and nsp8 to efficiently bind RNA and gain processivity *in vitro*, suggesting that together nsp12, nsp7, nsp8 are the minimal components that constitute the viral RNA polymerase (Fig. 1) (11–13). Additional functions are present in other nsps, such as helicase, methyltransferase, endonuclease, and transmembrane domains (7, 14). Some of these functions have been proposed to contribute to using the positive-sense RNA genome as a template to produce a full-length negative-sense genome RNA replication intermediate, as well as negative-sense subgenome RNAs that served as template for subgenomic mRNA synthesis (7, 15). The latter RNAs encode the structural and accessory proteins required to make new virus particles, such as the spike (S; RNA 2) and nucleocapsid (N; RNA 9) proteins (Fig. 1A).

**Figure 1.**
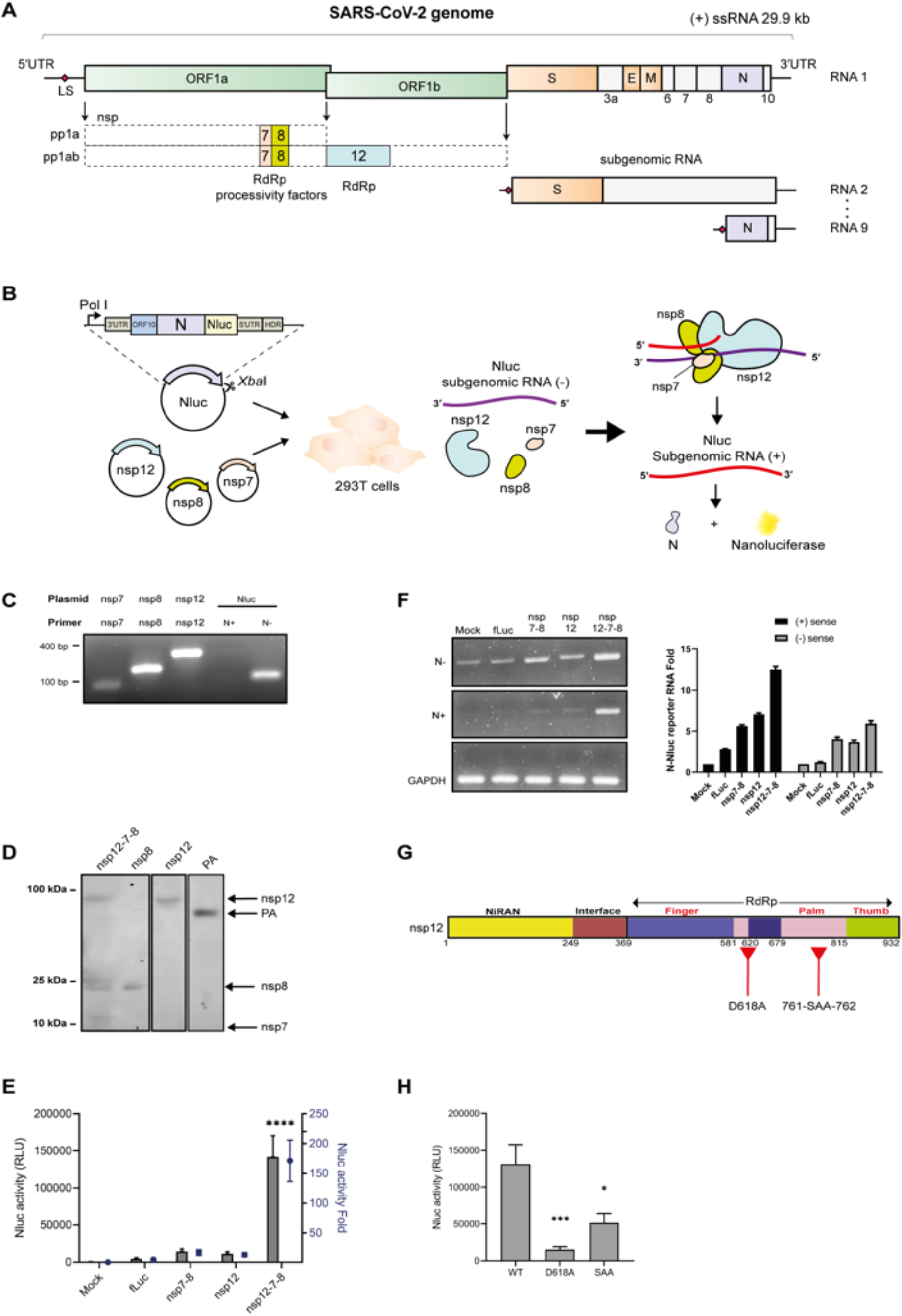
Design of the SARS-CoV-2 mini-genome assay. (**A**) Schematic diagram of SARS-CoV-2 genome showing the different open-reading frames (ORF), the location of nsp12, 7 and 8, as well as the subgenomic RNAs 2 and 9. (**B**) Schematic of the SARS-CoV-2 mini genome assay. Plasmids are transfected in HEK 293T cells to express SARS-CoV-2 nsp12, nsp7, and nsp8. A reporter plasmid containing a negative sense ORF encoding nanoluciferase (Nluc) inserted upstream of the SARS-CoV-2 N ORF. The untranslated regions (UTR) of the SARS-CoV-2 genome and ORF10 are also present in the reporter plasmid. Expression of the reporter plasmid yields a negative-sense RNA. The SARS-CoV-2 polymerase complex copies the negative-sense RNA into a positive-sense reporter RNA that can be translated by the host cell machinery to produce the Nluc and N proteins. (**C**) RT-PCR analysis of nsp7-flag, nsp8-flag, nsp12-flag and negative-sense Nluc-N gene expression 24 hours after transfection. (**D**) Expression of SARS-CoV-2 nsp7-flag, nsp8-flag and nsp12-flag 24 hours after transfection. A pcDNA3-PA-flag plasmid encoding the flag-tagged influenza A virus polymerase PA subunit was used as expression control. (**E**) Nluc activity 24 h after nsps and reporter plasmid transfection. (**F**) Expression of negative-sense (-) and positive-sense (+) reporter RNAs as determined by RT-PCR (left panel) and RT-qPCR (right panel) 24 hours after transfection. GAPDH mRNA was detected as loading control. (**G**) Schematic of functional nsp12 domains and motifs. The location of nsp12 RdRp motifs A and motif C is indicated. (**H**) Effect of nsp12 mutations D618A and SDD=>SAA on Nluc reporter activity following transfection of nsp12, nsp7, nsp8. Error bars represent three biological repeats.

The key role of the viral RNA polymerase in the viral replication makes it an important target for antiviral drug development. Various compounds are currently being tested as treatment for COVID-19, with remdesivir showing inhibition of SARS-CoV-2 RNA synthesis *in vitro* and *in vivo* (16–18). However, the ability of other drugs to reduce virus spread or COVID-19 mortality in humans has been limited (19, 20). Moreover, the emergence of novel SARS-CoV-2 variants, containing mutations and deletions in the viral spike protein (4, 5, 21, 22) as well as the RNA polymerase subunits (23, 24), and the slow increase in immune pressure with the world-wide roll-out of vaccines, suggest that variants may arise that can escape current vaccine treatments. These points underline the importance of continued and improved antiviral research efforts and better methods to investigate the function of emerging mutations.

Currently, antiviral screens and mutational analyses of SARS-CoV-2 RNA synthesis are chiefly performed by labs with access to a BSL3 facility or experience with the manipulation of long RNA replicons (25–29). This limits the effort to analyze emerging adaptive mutations and the development of a better understanding of SARS-CoV-2 transcription and replication. To make SARS-CoV-2 replication analysis simpler, we developed a SARS-CoV-2 mini-genome assay that can be performed in a normal laboratory setting with four small plasmids that can be easily manipulated. The assay involves the expression of the minimal RNA polymerase complex composed of the nsp7, nsp8 and nsp12, alongside a negative-sense RNA reporter encoding a fusion between the SARS-CoV-2 nanoluciferase (Nluc) and the N protein. The activity of nsp12 converts the negative sense reporter into positive-sense RNA that can be translated. This activity is sensitive to mutations in the RdRp domain of nsp12 as well as treatment with remdesivir. Overall, our approach facilitates measurements with lower background signals for increased sensitivity and allows straightforward construction of polymerase complexes of different SARS-CoV-2 VOC or specific mutations by side-directed mutagenesis for fundamental studies of SARS-CoV-2 replication.

## RESULTS

### Construction and validation of the SARS-CoV-2 mini genome assay

The SARS-CoV-2 positive-sense genome is ^~^30 kb (Fig. 1A), which complicates cloning and *in vitro* transcription. Moreover, often these long constructs are often not stable, need to be split in two or more parts for biosafety reasons, or require electroporation for transient expression (26, 27, 29). Shorter SARS-CoV-2 replicons have recently been reported that involve expression of a positive-sense RNA, containing the 5’ and 3’ UTRs of the genome and an internal reporter gene, which is amplified by individual nsps or polyproteins expressed from additional plasmids (25, 30). However, the expression of a positive-sense reporter RNA raises the background of the assay, because the positive-sense viral RNA can function as regular messenger RNA, reducing the sensitivity and range of these experiments.

To develop a simple read-out of SARS-CoV-2 RNA synthesis with better sensitivity, we used the fact that SARS-CoV-2 expresses its genome through a nested set of negative-sense subgenomic RNAs (7). These negative-sense subgenomic RNAs are subsequently transcribed into a nested set of positive-sense viral mRNAs by the viral RNA polymerase (Fig. 1A) (7, 15). Taking the second-smallest negative-sense subgenomic RNA (encoding for the viral N protein; Fig. 1A) as basis, we next inserted a negative-sense open reading frame (ORF) encoding Nluc upstream of the ORF encoding N (Fig. 1B), allowing stop-and-go translation of the two ORFs without additional regulatory elements that might affect SARS-CoV-2 replication. The 5’ and 3’ untranslated regions (UTR) that flank the ORF and contain the regulatory elements needed for the transcription of negative-sense SARS-CoV-2 subgenomic RNAs were left untouched. A 10-nt long polyU tail was added to the 5’ as template, in line with recent findings (31). The total length of this recombinant subgenomic RNA is 2471 nt, including polyU-tail (see supplemental information for sequence). In essence, this construct is similar to a positive-sense defective intermediate that was previously used in combination with a helper coronavirus to study CoV RNA synthesis (32).

To drive the expression of the negative-sense SARS-CoV-2 Nluc-N reporter, we inserted the Nluc-N reporter downstream of a cellular RNA polymerase I (Pol I) promoter sequence (Fig. 1B) that has been successfully used to for the expression of influenza A virus negative-sense RNA molecules (33). A hepatitis delta virus ribozyme (HDR) was inserted downstream of the 3’ terminus of the negative-sense Nluc-N reporter, to ensure formation of a native 3’ terminal sequence. Transfection of the reporter plasmid (Fig. 1B; see supplemental information for sequence) into HEK 293T cells resulted in the production of negative-sense RNA, but no positive-sense RNA (Fig. 1C).

Structural and biochemical experiments have shown that SARS-CoV-2 nsp12 depends on nsp7 and nsp8 for efficient RNA binding and processivity (9), and recent single-molecule experiments show that this complex can copy at least 1043 nt without the need for additional accessory nsps (34). To achieve transcription of the negative-sense Nluc-N reporter to positive-sense RNA, we co-transfected plasmids expressing flag-tagged nsp7, nsp8, and nsp12 in a 1:1:1 ratio to form the minimal complex required for polymerase activity. We confirmed expression of nsp-encoding genes by RT-PCR of nsp mRNAs and western blot analysis (Fig. 1C, D). To obtain replication of our negative-sense reporter RNA, we transfected the plasmids encoding nsp12, nsp8 and nsp7, and the Nluc-N reporter into HEK 293T cells. After 24 h, cells were harvested and Nluc activity was measured. In the presence of nsp7, nsp8 and nsp12, an Nluc signal was measured that was on average 170-fold higher than background following several rounds of optimization (Fig. 1E). We found that a ratio of 8:8:8:1 for the three nsps and the Nluc-N reporter was optimal for our plasmids. In addition, we observed a well-defined increase in both the negative-sense and positive-sense Nluc-N RNA signal in these reactions as measured by RT-qPCR (Fig. 1E, F). No significant increase in the Nluc signal was measured when the Nluc-N reporter was expressed with nsp12, or nsp8 and nsp7 alone, although a minor increase in the negative-sense and positive-sense RNA signals was observed in the presence of these proteins (Fig. 1F). Different nsp12, nsp8 and nsp7, and Nluc-N reporter ratios yielded lower Nluc signals in our hands (Fig. 2). Addition of empty plasmid to our assay only had minor impact on the signal at amounts 5 times higher than our standard input (Fig. S1).

**Figure 2.**
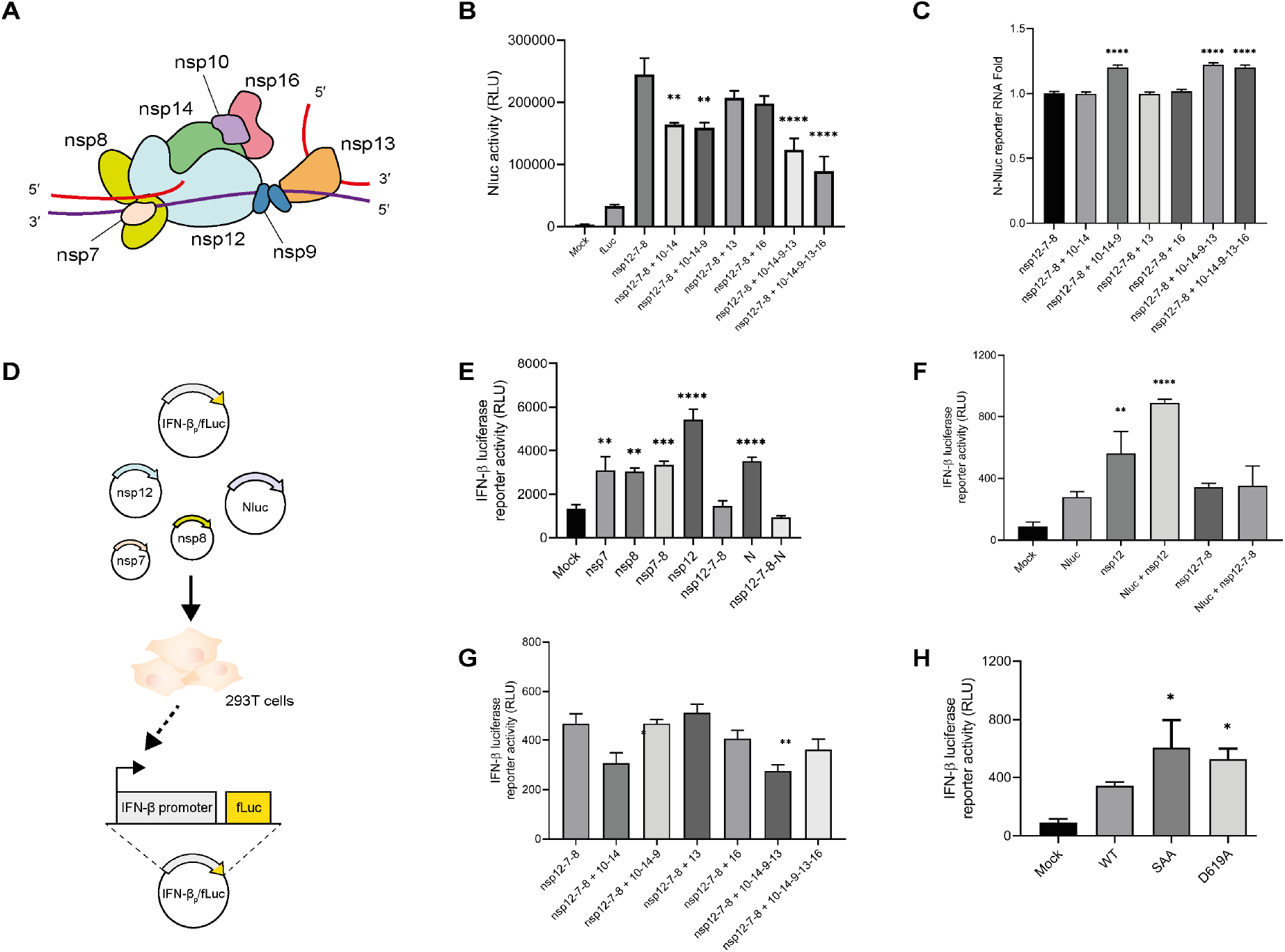
Impact of nsp complex formation on innate immune activation. (**A**) Model of the SARS-CoV-2 replication-transcription complex (RTC). The RTC core is composed by nsp12, nsp7 and nps8. The exonuclease nsp14 and its co-factor nsp10 may convey proofreading activity (38). Methylation of capped RNA likely performed by nsp16. Helicase nsp13 is likely involved in unwinding dsRNA structures and nsp9 binds single-stranded RNA, but may also play a role in capping (42, 44). (**B**) Impact of co-expression of other SARS-CoV-2 nsps on Nluc-N reporter activity 24h after transfection. Error bars show six biological repeats from two independent experiments. For statistical testing, a group comparison analysis was performed against the nsp12-nsp7-nsp8 combination. (**C**) Expression of positive-sense reporter RNAs as determined by RT-qPCR 24 hours after transfection. Error bars show three biological repeats. For statistical testing, a group comparison analysis was performed against the nsp12-nsp7-nsp8 combination. (**D**) Experimental design of the firefly luciferase-based IFN expression assay. HEK293T cells are co-transfected with a firefly luciferase reporter plasmid under the control of the IFN-β promoter (pIFΔ(−116)lucter) and different SARS-CoV-2 plasmids. (**E**) IFN-β promoter activity after expression of SARS-CoV-2 nsp7, nsp8, nsp12 and N protein. (**F**) IFN-β promoter activity after expression of the Nluc-N reporter, nsp12 and nsp12-7-8 for the minimal replication complex. (**G**) IFN-β promoter activity after expression of the Nluc-N reporter and others nsp involved in the large replication complex. For statistical testing, a group comparison analysis was performed against the nsp12-nsp7-nsp8 combination. (**H**) IFN-β promoter activity following expression of wild-type or mutant nsp12 in the presence of the nsp12-nsp7-nsp8 complex. In figures E-H, six biological replicates were collected from two independent experiments. Error bars show SEM. (E, F, H) For statistical testing, a group comparison analysis was performed against the mock condition. *P < 0.05, **P < 0.01; ***P < 0.001 ****P < 0.0001, as determined by Dunnett’s multiple comparisons test.

To further confirm that the measured Nluc signal was dependent on the activity of the minimal SARS-Cov-2 RNA polymerase complex, we mutated the conserved aspartates of motif A (D619) and C (D761 and D762) of the nsp12 RdRp domain (Fig. 1G) to alanine, creating mutants D619A and SAA, respectively. It has previously been shown that motifs motif A (D619) and C (D761 and D762) are necessary for RNA polymerase activity (8, 35–37). Co-transfection of these nsp12 mutants alongside wildtype nsp7 and nsp8, significantly reduced the NLuc signal, as well as the negative-sense and positive-sense RNA levels (Fig. 1H), in line with the negative effect of these mutations on nsp12 activity in *in vitro* experiments (8, 35–37). Together, these results suggest that nsp7, nsp8 and nsp12 are sufficient to replicate a negative-sense SARS-CoV-2 subgenomic RNA in cell culture. Because no full-length viral genome is used and all structural viral genes except N are omitted from the construct, no virus can be produced using this system, making it safe to use under standard laboratory conditions.

### Effect other nsps and induction of IFN-β promoter activity by individual nsps

The SARS-CoV-2 genome encodes many nsps that form a larger complex with nsp7, nsp8 and nsp12 (Fig. 2A), and affect the activity of the RNA polymerase in cell culture. For instance, nsp14 has been proposed to affect the error rate of CoV RNA synthesis due to its exonuclease activity, while nsp16 may affect capping and translation of the viral RNAs due to its methyltransferase activity (38). Interestingly, expression of other nsps alongside the minimal SARS-CoV-2 RNA polymerase subunits nsp7, nsp8 and nsp12 reduced Nluc activity by 1.5-fold for the expression of nsp10 and nsp14, and 2.7-fold for the expression of nsp9, nsp10, nsp13, nsp14 and nsp16 (Fig. 2B). However, positive-sense RNA expression was slightly increased when nsp10, nsp14 and nsp9 were co-expressed with the nsp12/7/8 complex (Fig. 2C). This discrepancy between higher viral mRNA levels and lower Nluc activity suggests that less mRNA may be available for translation and that nsp9, nsp10, nsp14 contribution to the formation or stability of a replication complex. Plasmid mRNA expression for each nsps was confirmed 24h after co-transfection (Fig. S2A-B).

To investigate if the absence of other nsps from the nsp12/7/8 complex led to more aberrant RNA or dsRNA synthesis, innate immune signaling and a subsequent activation of IFN-β promoter activity, we co-transfected a plasmid encoding firefly luciferase downstream of the IFN-β promoter (Fig. 2D). As control conditions, we transfected the individual expression plasmids alone. Interestingly, expression of N, nsp7, nsp8 or nsp12 alone, or a combination of nsp7 and 8, let to a significant increase in IFN-β promoter activity (Fig. 2E). By contrast, expression of the three RNA polymerase subunits together did not result in a significant induction of IFN-β promoter activity (Fig. 2E). Expression of the Nluc-N reporter RNA resulted in an increase of IFN-β promoter activity relative to our mock transfection (Fig. 2F). Co-transfection of nsp12 alongside the Nluc-N reporter raised the IFN-β promoter activity further (Fig. 2F), while expression of the minimal RNA polymerase did not raise the IFN-β promoter activity relative to the Nluc-N reporter alone. Moreover, co-expression of other nsps (i.e., nsp10, nsp14, nsp9, nsp13, nsp16) associated with the nsp12/7/8 complex and the Nluc-N reporter RNA, did not increase the IFN-β promoter activity found with the minimal replication complex (Fig. 2G). By contrast, co-expression of nsp10 and nsp14, or nsp10, nsp14, nsp9 and nsp13 decreased the IFN-β promoter activity, suggesting that these larger complexes may better shield the reporter RNA from host pathogen receptors than the nsp12/7/8 complex. To investigate if nsp12 polymerase activity had an effect of on the IFN-β promoter activity, we transfected nsp12 wt or active site mutants D619A and SAA together with nsp7, nsp8 into HEK 293T cells (Fig. 2H). Interestingly, the wt minimal RNA polymerase complex induced less IFN-β promoter activity than the two active site mutants.

Together, these observations suggest that the minimal RNA polymerase complex consisting of nsp7, nsp8 and nsp12 forms a stable, active complex that is sufficient for the replication of a subgenomic RNA. In addition, these findings suggest that the complex is less prone to detection by the host cell than the subunits alone. The observations also suggest that different nsp constellations have different replication/transcription activities, and that the nsp constellation affects the dynamics of viral RNA synthesis and mRNA translation.

### Inhibition of mini-genome assay by antiviral compounds

To investigate whether the activity of the minimal SARS-CoV-2 RNA polymerase complex can be inhibited by antivirals in cell culture, we next treated transfected cells 3 hours post transfection (Fig. 3A) with different concentrations of the prodrugs remdesivir, ribavirin, favipiravir (also known as T-705), 5-fluorouracil (also known as 5-FU), or a DMSO control. Twenty-four hours after the addition of the compounds, the cells were lysed and the cytotoxicity and Nluc levels measured (Fig. 3A). Remdesivir, ribavirin, and favipiravir did not show significant cytoxocity over the compound concentration range tested (Fig. 3B). By contrast, 5-fluorouracil showed high levels of cytotoxicity and was not further considered. Analysis of the effect on SARS-CoV-2 RNA polymerase activity revealed a strong inhibitory effect remdesivir on the Nluc activity level (IC_50_ 6.3 μM; Fig. 3B). This observation is in line with the remdesivir IC_50_ observed in virus infections and other replicon assays (25, 26). To confirm that the reduction in Nluc activity level was correlated with a reduction in the positive-sense Nluc-N reporter RNA level, we performed an RT-qPCR analysis on the remdesivir-treated cells. As shown in Fig. 3C, we found that remdesivir also significantly reduced the Nluc-N RNA level, although at a higher concentrations than observed for the Nluc signal. The compounds ribavirin and favipiravir modestly reduced the Nluc levels, but only at higher concentrations (IC_50_ >500 μM; Fig. 3B). Reports on the effect of T-705 and ribavirin on SARS-CoV-2 largely indicate that the effect is weaker than that of remdesivir (13, 39–41), in line with our observations. Overall, these results indicate that the subgenome RNA-based reporter assay can be used as approximation of SARS-CoV-2 RNA synthesis and used to measure the inhibitory potential of antivirals.

**Figure 3.**
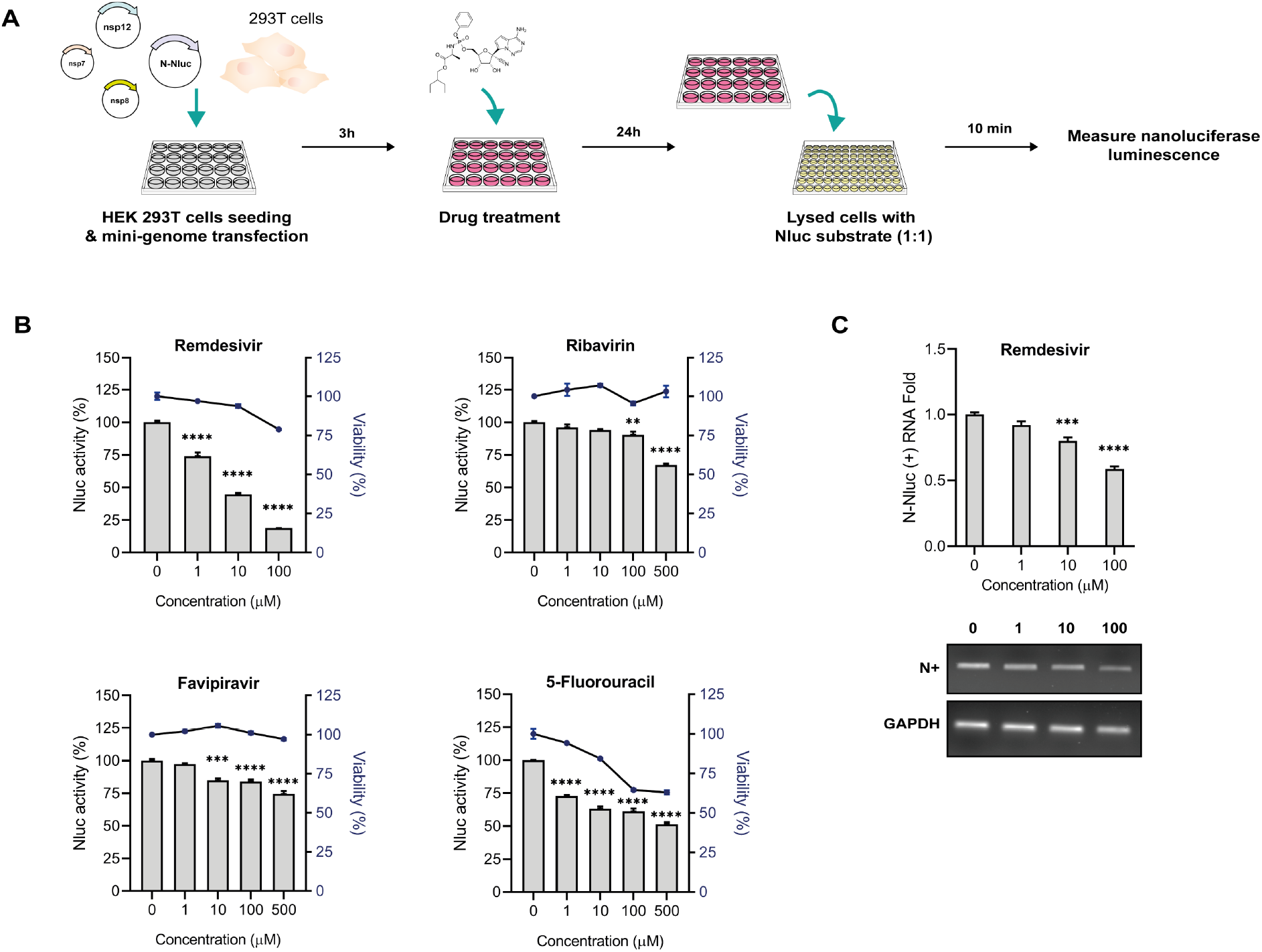
Inhibition of nanoluciferase activity by antiviral compounds. (**A**) Experimental design of the mini-genome inhibition assay with antivirals. Approximately 1×10^5^ HEK 293T cells suspended in Optimem were co-transfected with plasmids expressing Nluc-N reporter, nsp7-myc, nsp8-myc, and nsp12-myc in a 1:8:8:8 ratio and subsequently seeded in P24-well plate. Three hours after transfection, antiviral compounds were added to the media and incubated on the cells for 24 hours. Cells were finally lysed and incubated with nanoluciferase substrate for 10 min prior to measurement of the luminescence signal. (**B**) Relative nanoluciferase activity after exposure to antiviral compounds. Remdesivir, ribavirin, favipiravir or 5-fluorouracil were added to cell media 3 h post-transfection at the concentrations indicated. Nanoluciferase activity and cytotoxicity were measured 24 h post-treatment. A final concentration of 0.5% DMSO was used as 0 μM antiviral concentration. Each bar represents the average of six biological replicates that were collected over two independent experiments. (**C**) Positive-sense Nluc-N expression after remdesivir treatment. Total RNA was extracted from cells treated with remdesivir and retrotranscribed. qPCR and PCR analyses were performed for the N-encoding positive-sense sequence. GAPDH mRNA was used as internal control. The bars in the qPCR analysis represent three biological repeats. Error bars show SEM. **P < 0.01; ***P < 0.001, ***P < 0.0001 as determined by Dunnett’s multiple comparisons test.

### Effect of emerging nsp mutations on RNA polymerase activity

Since the emergence of SARS-CoV-2 in 2019, several new strains have appeared that contain mutations in the spike protein that improve SARS-CoV-2 spread or receptor binding (4, 5, 21, 22, 24). Amino acid substitutions in VOC have also appeared in the nsps that make up the core of the RNA polymerase (Fig. 4A) (23) and their location in the structure suggest that they may affect the interaction between the nsps in the complex, potentially affecting viral RNA synthesis (Fig. 4B, C). To investigate whether these mutations alone or in combination affect the activity of the minimal SARS-CoV-2 RNA polymerase complex, we introduced them into our plasmids encoding nsp7, nsp8 and nsp12, and expressed the mutant proteins in our mini-genome assay. Using stringent statistical testing, we find that none of the mutations significantly changed the activity of the RNA polymerase complex (Fig. 4D). However, a combination of mutations nsp12 P323L and nsp7 S25L negatively affected the activity of the RNA polymerase complex (Fig. 4D). RT-qPCR analysis showed that positive-strand RNA expression was also slightly reduced for this mutation combination (Fig. 4E). These observations are in line with a previous analysis of the RNA polymerase structure suggesting that the above point mutations may affect protein-protein interaction interfaces in the complex and potentially the activity of the RNA polymerase (Fig. 4B, C).

**Figure 4.**
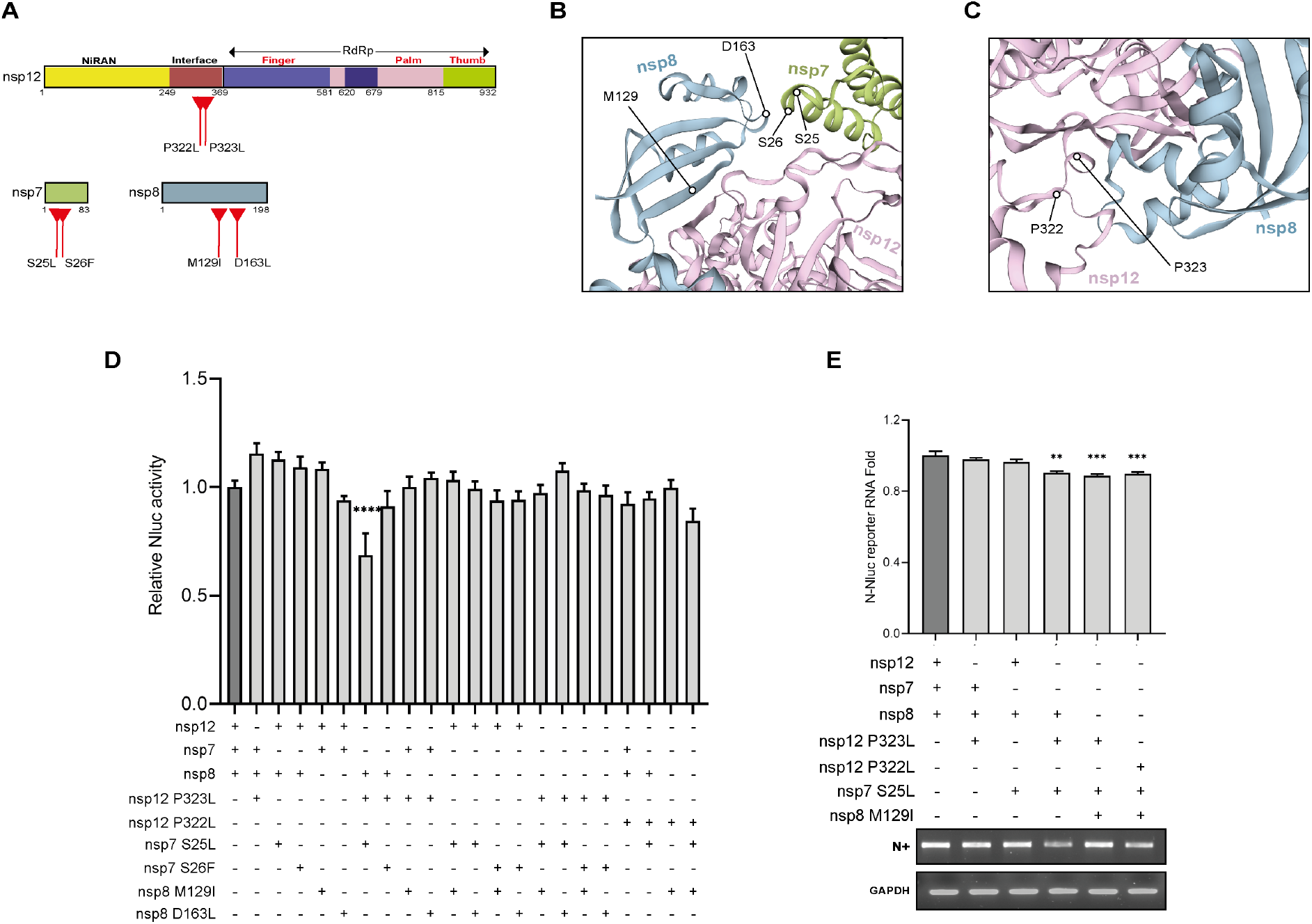
Impact of adaptive mutation on the activity of the nsp7-nsp8-nsp12 complex. (**A**) Schematic of functional domains and motifs of nsp7, nsp8 and nsp12. The location of the nsp7, nsp8 and nsp12 mutations is indicated. (**B**) Close-up of the SARS-CoV-2 RNA polymerase structure based on modelling described in supplemental text, with nsp12 in pink, nsp7 in green, and nsp8 in blue. Location of nsp7 and nsp8 is shown. (**C**) Location of nsp12 mutations in SARS-CoV-RNA polymerase structure. (**D**) Effect of nsp12 mutations P323L and P322L, nsp7 mutations S25L and S26F, nsp8 mutations M129I and D163L on Nluc reporter activity. Bars represent mean of six biological repeats from two independent experiments. Error bars indicate SEM. (**E**) Expression of positive-sense reporter RNAs as determined by RT-qPCR (upper panel) and RT-PCR (lower panel) 24 hours after transfection. GAPDH mRNA was detected as loading control. Bars represent mean of three biological repeats. Error bars indicate SEM. **P < 0.01; ***P < 0.001, ***P < 0.0001 as determined by Dunnett’s multiple comparisons test.

## Discussion and conclusions

Despite the successful rapid development of vaccines against SARS-CoV-2, there is still an immediate need for antivirals against current and future SARS-CoV variants. Antiviral screens and mutational analyses of SARS-CoV-2 RNA synthesis are currently chiefly performed by labs with access to a BSL3 facility or experience with the manipulation of long RNA replicons (25–29). This slows down world-wide efforts to rapidly analyze emerging adaptive mutations, screening of potential drugs identified through modeling, and the development of a better understanding of SARS-CoV-2 transcription and replication. To make the analysis of SARS-CoV-2 replication simpler and more widely available, we developed a SARS-CoV-2 mini-genome assay that can be performed in a normal tissue culture setting and that produces a signal that is on average 170-fold higher than background. This sensitivity is well above recently reported signals obtained with other replicons (30). This improved sensitivity likely derives from the fact that we use a negative-sense reporter RNA, instead of a positive-sense reporter RNA. The results of our assay also show that the minimal SARS-CoV-2 RNA polymerase complex, composed of the nsp7, nsp8 and nsp12, is sufficient to replicate a 2.5 kb subgenomic RNA and create positive-sense RNA products that can be translated.

The efficient replication of a negative sense subgenomic RNA in cell culture and the production of translation-competent positive sense RNAs by the minimal RNA polymerase complex, raises a plethora of questions about the role of the other nsps during SARS-CoV-2 replication and our current understanding of CoV RNA synthesis in general. Our observations that the combination of nsp7, nsp8 and nsp12 is, at the least, modestly processive is supported by biochemical, single-molecule and other replicon studies (30, 34, 36), suggesting that the role of the nsp13 helicase may be more essential for a longer RNA template or subgenomic RNA synthesis (42). SARS-CoV-2 nsp14 and nsp10 also appear to be accessory factors, in line with their potential role as proofreading enzymes (30, 36) or in innate immune evasion (43), but we do observe that their presence stimulates viral replication and likely reduces viral mRNA translation. Also remarkable is the production of translation-competent subgenomic RNAs by nsp12, nsp7 and nsp8 alone, since capping of SARS-CoV-2 RNAs has been proposed to involve nsp9, nsp12, nsp14 and nsp16 (44, 45). Finally, it remains unresolved how nsp12, which contains a primer-dependent RdRp domain without priming-loop (8, 9), is able to perform *de novo* initiation in the presence of nsp7 and nsp8. Previous reports suggest that nsp8 or a complex nsp7 and nsp8 can synthesize short RNA primers (46, 47), but a biochemical understanding of how these activities are coupled is currently lacking. We will use our assay and modifications of it to address these and other questions in the future.

In conclusion, the described method facilitates sensitive measurements of SARS-CoV-2 RNA polymerase activity with low background signals. We demonstrate that this method can be used to study the RNA synthesis activity of SARS-CoV-2 variants or specific RNA polymerase mutations, screen for novel anti-vital compounds or perform fundamental studies of SARS-CoV replication. We hope that we and others can use the principle of our assay to identify novel anti-SARS-CoV-2 drugs and to study these fundamental questions in more detail in the future.

## MATERIALS AND METHODS

### Cells

Human embryonic kidney (HEK) 293T cells (ATCC CRL-3216) were cultured in Dulbecco’s Modified Eagle Medium (DMEM) (PAN-Biotech) with 10% feral calf serum (FCS) (Sigma-Aldrich), 1% L-Glutamine (Sigma-Aldrich). Cells were maintained in vented culture flasks in a humidified incubator at 5% CO_2_ and 37°C.

### Plasmids

Plasmids pcDNA6.B-nsp7-flag, pcDNA6.B-nsp8-flag, pcDNA6.B-nsp12-flag, pcDNA6.B-nsp10-flag, pcDNA6.B-nsp14-flag, pcDNA6.B-nsp9-flag, pcDNA6.B-nsp13-flag, pcDNA6.B-nsp16-flag, pcDNA6.B-N-flag containing codon-optimized versions of SARS-CoV-2 strain Wuhan-Hu-1 (2019) nsp7, nsp8, nsp12, nsp10, nsp14, nsp9, nsp13, nsp16 and nucleocapsid respectively, were used for the expression of nsp7, nsp8, nsp12, nsp10, nsp14, nsp9, nsp13, nsp16 and nucleocapsid. Mutations in nsp12 (motif A (D619A), motif C (DD761-762AA), P323L, P322L), in nsp7 (S25L, S26F), and in nsp8 (M129I, D163L) were introduced by site-directed mutagenesis of pcDNA6.B-nsp12-flag, pcDNA6.B-nsp7-flag or pcDNA6.B-nsp8-flag using primers listed in Table S1. Mutations were confirmed by Sanger sequencing.

To construct the negative-sense RNA reporter, we designed construct PolI-SARS-CoV2-NLuc-N-HDR and ordered this as insert in the pMK-RQ backbone (Invitrogen construct ID 20ACX40P). The resulting plasmid pPolI-SARS-CoV2-NLuc-N was digested with XbaI prior to transfection to eliminate background (read-through) transcription from other genes encoded in the reporter plasmid. Transfection control pcDNA3-FF-luciferase and IFN-β reporter plasmid pIFΔ(−116)lucter were described previously (48, 49). All plasmids were amplified in Turbo cells (NEB) and isolated using a high purity plasmid prep kit (Thermo Fisher). Plasmid inserts were confirmed by double digestion with BamHI/SacII for nsp constructs and with XbaI/StuI for the N-NLuc reporter construct (Fig. S2A).

### Mini genome assay

To measure SARS-CoV-2 polymerase activity in cell culture, approximately 2×10^5^ 293T cells were transfected with 200 ng each of pcDNA6.B-nsp7-flag, pcDNA6.B-nsp8-flag, and pcDNA6.B-nsp12-flag, and 25 ng linear pPolI-SARS-CoV2-NLuc-N plasmid using 2μl of Lipofectamine 2000 (Invitrogen) to a DNA:lipofectamine ratio of 1:3 and 100μl Opti-MEM (Gibco). Cells were collected 24 hours post-transfection, washed with ice-cold PBS, and split into equal fractions for separate luminescence, western and RNA analysis. For western analysis, the cells were lysed in cold RIPA buffer (50mM Tris pH 7.8, 150mM NaCl, 1% NP-40, 0.5% sodium deoxycholate, 0.1% SDS, 1 mM DTT and 1X protease inhibitor cocktail (Sigma-Aldrich). For RNA analysis, the lysed cells were suspended in TRIzol (Invitrogen) and total RNA extracted as described previously (49). For firefly luciferase and Nluc activity assays, cells were analyzed using a Nano-Glo Dual-Luciferase Reporter Assay (Catalogue number N1610, Promega) kit according to the manufacturer’s protocol and GloMax Navigator Microplate Luminometer (Promega). For each experiment shown, at least three biological replicates were conducted.

### Western blot

HEK 293T cells lysed in RIPA buffer were suspended in SDS loading buffer (180 mM Tris pH 6.8, 4% SDS, 0.2% bromophenol blue, 20% glycerol, 0.4 M DTT) and denatured for 5 min at 95°C. Proteins were separated by SDS-PAGE using 4-20% Mini-PROTEAN TGX Precast gel (Bio-Rad) and transferred to a nitrocellulose membrane (GVS) using a Trans-Blot Turbo transfer system (Bio-Rad). Membranes were incubated with blocking buffer (PBS, 0.05% Tween-20, 10% milk) for 1h, and subsequently followed by primary antibody diluted in blocking buffer overnight at 4°C. The nsp7, nsp8 and nsp12 proteins were subsequently detected using the mouse monoclonal anti-DYKDDDDK M2 antibody (Catalogue number F3165, Sigma-Aldrich) diluted 1:1000 in TBST/5% milk. Cellular proteins were detected using rabbit polyclonal antibodies anti-GAPDH (Catalogue number GTX100118, GeneTex) diluted 1:4000 in TBST/5% milk. The SARS-CoV-2 N protein was detected using a monoclonal rabbit antibody (Catalogue number 40143-R019, SinoBiological) diluted 1:1000 in TBST/5% milk. Secondary antibodies IRDye 680 goat anti-mouse (Catalogue number 926-68020, Li-cor) and IRDye 800 donkey antirabbit (Catalogue number 926-32213, Li-cor) were used to detect western signals with an Odyssey scanner (Li-cor).

### Luciferase-based IFN expression assay

HEK 293T cells were transfected with SARS-CoV-2 nsp plasmids as described above for the mini-genome assay and co-transfected with 100 ng of a firefly luciferase reporter plasmid under the control of the IFN-β promoter as described previously (48). The luciferase expression levels were measured 24h post-transfection using a Nano-Glo Dual-Luciferase Reporter Assay kit (Catalogue number N1610, Promega) according to the manufacturer’s protocol.

### Cell viability

The HEK 293T cell viability in the presence of compounds was measured using a CellTiter-Glo 2.0 (Catalogue number G9241, Promega) based on the manufacturer’s protocol. Compounds were added to DMEM containing 10% FCS and 1% L-glutamine and incubated for 24h.

### RNA extraction, PCR and RT-qPCR

Total RNA from 293T cells was isolated using TRIzol (Invitrogen) as described previously. To make cDNA of negative-sense RNAs, we used random hexamers in combination with Superscript III reverse transcriptase (Invitrogen) as described previously (48). To make cDNA of positive-sense RNAs and cellular mRNAs, we used oligodT_18_ in combination with Superscript III reverse transcriptase (Invitrogen). PCRs were performed using Taq DNA polymerase (Qiagen) and the primers listed in Table S2. Real-time quantitative PCR was carried out using the qPCR Brilliant III SYBR Master Mix (Agilent), the primers listed in Table S2, and a StepOnePlus Realtime PCR System (Applied Biosystems). Human glyceraldehyde-3-phosphate dehydrogenase (GAPDH) mRNA levels were measured as cellular control.

### Compound screening and IC_50_ measurement

Compounds were added into the growing medium with an initial concentration of 500 μM and further dilution. Remdevisir (catalogue number GS-5734, Tocris), ribavirin (catalogue number R9644, Sigma-Aldrich), favipiravir (catalogue number S79755MG, Selleck Chemical LLC) and 5-Fluorouracil (catalogue number F6627, Sigma-Aldrich) were prepared in DMSO. Cells were seeded and transfected in 24-well plates 3 h prior to compound addition to the medium. DMEM containing 0.5% DMSO was used as negative control. The Nluc assay was performed 24h post-transfection. A detailed protocol is described in the supplemental information.

### Protein structure modelling

Protein modelling was carried out with SWISS-MODEL. The wt and mutant nsp12-nsp7-nsp8 complexes were modelled with sequences of nsp12, nsp7 and nsp8 proteins based on NC_045512.2 reference sequence, SARS-CoV-2 strain Wuhan-Hu-1 (2019) (Table S3).

### Statistical analysis

Statistical analysis was performed with GraphPad PRISM 8 software. Values are reported as the mean +/- standard error of the mean (sem) of three biological replicates. Data were analysed with one-way Anova followed by Dunnett’s multiple comparison test or t-test. Two-tailed p-values lower than 0.05 were considered as statistically significant.

## Acknowledgments

The authors would like to thank M. van Hoesel for experimental support, and J. Edgar and W. Pei-Hui for plasmids.

