## Supplemental information for "A SARS-CoV-2 mini-genome assay based on negative-sense RNA to study replication inhibitors and emerging mutations"

### A simple SARS-coronavirus 2 mini-genome assay launched from negative-sense RNA

### Replicon sequence

#### Full replicon sequence:

TTTTTTTTTTGTCATTCTCCTAAGAAGCTATTAAAATCACATGGGGATAGCACTACTAAAATTAATTTTACACATTAGGGCTCTTC
CATATAGGCAGCTCTCCCTAGCATTGTTCACTGTACACTCGATCGTACTCCGCGTGGCCTCGGTGAAAATGTGGTGGCTCTTTC
AAGTCCTCCCTAATGTTACACACTGATTAAAGATTGCTATGTGAGATTAAAGTTAACTACATCTACTTGTGCTATGTAGTTACGA
GAATTCATTCTGCACAAGAGTAGACTATATATCGTAAACGGAAAAGCGAAAACGTTTATATAGCCCATCTGCCTTGTGTGGTCT
GCATGAGTTTAGGCCTGAGTTGAGTCAGCACTGCTCATGGATTGTTGCAATTGTTTGGAGAAATCATCCAAATCTGCAGCAGG
AAGAAGAGTCACAGTTTGCTGTTTCTTCTGTCTCTGCGGTAAGGCTTGAGTTTCATCAGCCTTCTTCTTTTTGTCCTTTTTAGGCT
CTGTTGGTGGGAATGTTTTGTATGCGTCAATATGCTTATTCAGCAAAATGACTTGATCTTTGAAATTTGGATCTTTGTCATCCAA
TTTGATGGCACCTGTGTAGGTCAACCACGTTCCCGAAGGTGTGACTTCCATGCCAATGCGCGACATTCCGAAGAACGCTGAAG
CGCTGGGGGCAAATTGTGCAATTTGCGGCCAATGTTTGTAATCAGTTCCTTGTCTGATTAGTTCCTGGTCCCCAAAATTTCCTTG
GGTTTGTTCTGGACCACGTCTGCCGAAAGCTTGTGTTACATTGTATGCTTTAGTGGCAGTACGTTTTTGCCGAGGCTTCTTAGA
AGCCTCAGCAGCAGATTTCTTAGTGACAGTTTGGCCTTGTTGTTGTTGGCCTTTACCAGACATTTTGCTCTCAAGCTGGTTCAAT
CTGTCAAGCAGCAGCAAAGCAAGAGCAGCATCACCGCCATTGCCAGCCATTCTAGCAGGAGAAGTTCCCCTACTGCTGCCTGG
AGTTGAATTTCTTGAACTGTTGCGACTACGTGATGAGGAACGAGAAGAGGCTTGACTGCCGCCTCTGCTCCCTTCTGCGTAGA
AGCCTTTTGGCAATGTTGTTCCTTGAGGAAGTTGTAGCACGATTGCAGCATTGTTAGCAGGATTGCGGGTGCCAATGTGATCTT
TTGGTGTATTCAAGGCTCCCTCAGTTGCAACCCATATGATGCCGTCTTTGTTAGCACCATAGGGAAGTCCAGCTTCTGGCCCAG
TTCCTAGGTAGTAGAAATACCATCTTGGACTGAGATCTTTCATTTTACCGTCACCACCACGAATTCGTCTGGTAGCTCTTCGGTA
GTAGCCAATTTGGTCATCTGGACTGCTATTGGTGTTAATTGGAACGCCTTGTCCTCGAGGGAATTTAAGGTCTTCCTTGCCATG
TTGAGTGAGAGCGGTGAACCAAGACGCAGTATTATTGGGTAAACCTTGGGGCCGACGTTGTTTTGATCGCGCCCCACTGCGTT
CTCCATTCTGGTTACTGCCAGTTGAATCTGAGGGTCCACCAAACGTAATGCGGGGTGCATTTCGCTGATTTTGGGGTCCATTAT
CAGATTACGCCAGAATGCGTTCGCACAGCCGCCAGCCGGTCACTCCGTTGATGGTTACTCGGAACAGCAGGGAGCCGTCGGG
GTTGATCAGGCGCTCGTCGATAATTTTGTTGCCGTTCCACAGGGTCCCTGTTACAGTGATCTTTTTGCCGTCGAACACGGCGAT
GCCTTCATACGGCCGTCCGAAATAGTCGATCATGTTCGGCGTAACCCCGTCGATTACCAGTGTGCCATAGTGCAGGATCACCTT
AAAGTGATGATCATCCACAGGGTACACCACCTTAAAAATTTTTTCGATCTGGCCCATTTGGTCGCCGCTCAGACCTTCATACGG
GATGATGACATGGATGTCGATCTTCAGCCCATTTTCACCGCTCAGGACAATCCTTTGGATCGGAGTTACGGACACCCCGAGATT
CTGAAACAAACTGGACACACCTCCCTGTTCAAGGACTTGGTCCAGGTTGTAGCCGGCTGTCTGTCGCCAGTCCCCAACGAAATC
TTCGAGTGTGAAGACCATTTTAGTTTGTTCGTTTAGAGAACAGATCTACAAGAGATCGAAAGTTGGTTGGTTTGTTACCTGGGA
AGGTATAAACCTTTAAT

#### Containing

3’ UTR:
TTTTTTTTTTTTGTCATTCTCCTAAGAAGCTATTAAAATCACATGGGGATAGCACTACTAAAATTAATTTTACACATTAGGGCTCT
TCCATATAGGCAGCTCTCCCTAGCATTGTTCACTGTACACTCGATCGTACTCCGCGTGGCCTCGGTGAAAATGTGGTGGCTCTTT
CAAGTCCTCCCTAATGTTACACACTGATTAAAGATTG

ORF10:
CTATGTGAGATTAAAGTTAACTACATCTACTTGTGCTATGTAGTTACGAGAATTCATTCTGCACAAGAGTAGACTATATATCGTA
AACGGAAAAGCGAAAACGTTTATATAGCCCAT

ORF9/nucleocapsid:
CTGCCTTGTGTGGTCTGCATGAGTTTAGGCCTGAGTTGAGTCAGCACTGCTCATGGATTGTTGCAATTGTTTGGAGAAATCATC
CAAATCTGCAGCAGGAAGAAGAGTCACAGTTTGCTGTTTCTTCTGTCTCTGCGGTAAGGCTTGAGTTTCATCAGCCTTCTTCTTT
TTGTCCTTTTTAGGCTCTGTTGGTGGGAATGTTTTGTATGCGTCAATATGCTTATTCAGCAAAATGACTTGATCTTTGAAATTTG
GATCTTTGTCATCCAATTTGATGGCACCTGTGTAGGTCAACCACGTTCCCGAAGGTGTGACTTCCATGCCAATGCGCGACATTC
CGAAGAACGCTGAAGCGCTGGGGGCAAATTGTGCAATTTGCGGCCAATGTTTGTAATCAGTTCCTTGTCTGATTAGTTCCTGGT
CCCCAAAATTTCCTTGGGTTTGTTCTGGACCACGTCTGCCGAAAGCTTGTGTTACATTGTATGCTTTAGTGGCAGTACGTTTTTG
CCGAGGCTTCTTAGAAGCCTCAGCAGCAGATTTCTTAGTGACAGTTTGGCCTTGTTGTTGTTGGCCTTTACCAGACATTTTGCTC
TCAAGCTGGTTCAATCTGTCAAGCAGCAGCAAAGCAAGAGCAGCATCACCGCCATTGCCAGCCATTCTAGCAGGAGAAGTTCC
CCTACTGCTGCCTGGAGTTGAATTTCTTGAACTGTTGCGACTACGTGATGAGGAACGAGAAGAGGCTTGACTGCCGCCTCTGC

TCCCTTCTGCGTAGAAGCCTTTTGGCAATGTTGTTCCTTGAGGAAGTTGTAGCACGATTGCAGCATTGTTAGCAGGATTGCGGG
TGCCAATGTGATCTTTTGGTGTATTCAAGGCTCCCTCAGTTGCAACCCATATGATGCCGTCTTTGTTAGCACCATAGGGAAGTCC
AGCTTCTGGCCCAGTTCCTAGGTAGTAGAAATACCATCTTGGACTGAGATCTTTCATTTTACCGTCACCACCACGAATTCGTCTG
GTAGCTCTTCGGTAGTAGCCAATTTGGTCATCTGGACTGCTATTGGTGTTAATTGGAACGCCTTGTCCTCGAGGGAATTTAAGG
TCTTCCTTGCCATGTTGAGTGAGAGCGGTGAACCAAGACGCAGTATTATTGGGTAAACCTTGGGGCCGACGTTGTTTTGATCG
CGCCCCACTGCGTTCTCCATTCTGGTTACTGCCAGTTGAATCTGAGGGTCCACCAAACGTAATGCGGGGTGCATTTCGCTGATT
TTGGGGTCCATTATCAGA

Nanoluc:
TTACGCCAGAATGCGTTCGCACAGCCGCCAGCCGGTCACTCCGTTGATGGTTACTCGGAACAGCAGGGAGCCGTCGGGGTTG
ATCAGGCGCTCGTCGATAATTTTGTTGCCGTTCCACAGGGTCCCTGTTACAGTGATCTTTTTGCCGTCGAACACGGCGATGCCTT
CATACGGCCGTCCGAAATAGTCGATCATGTTCGGCGTAACCCCGTCGATTACCAGTGTGCCATAGTGCAGGATCACCTTAAAGT
GATGATCATCCACAGGGTACACCACCTTAAAAATTTTTTCGATCTGGCCCATTTGGTCGCCGCTCAGACCTTCATACGGGATGA
TGACATGGATGTCGATCTTCAGCCCATTTTCACCGCTCAGGACAATCCTTTGGATCGGAGTTACGGACACCCCGAGATTCTGAA
ACAAACTGGACACACCTCCCTGTTCAAGGACTTGGTCCAGGTTGTAGCCGGCTGTCTGTCGCCAGTCCCCAACGAAATCTTCGA
GTGTGAAGACCAT

5’ UTR:
TTTAGTTTGTTCGTTTAGAGAACAGATCTACAAGAGATCGAAAGTTGGTTGGTTTGTTACCTGGGAAGGTATAAACCTTTAAT

#### Plasmid flanking sequences

Pol I promoter upstream of 3’ UTR:
ACGGGCCGGCCCCTGCTTGTGGCACGGGCGGCCGGGAGGGCGTCCCCGGCCCGGCGCTGCTCCCGCGTGTGTCCTGGGGTTG
ACCAGAGGGCCCCGGGCGCTCCGTGTGTGGCTGCGATGGTGGCGTTTTTGGGGACAGGTGTCCGTGTCGCGCGTCGCCTGGG
CCGGCGGCGTGGTCGGTGACGCGACCTCCCGGCCCCGGGGGAGGTATATCTTTCGCTCCGAGTCGGCATTTTGGGCCGCCGG
GTTATT

HDR downstream of 5’ UTR:
GGCCGGCATGGTCCCAGCCTCCTCGCTGGCGCCGGCTGGGCAACATTCCGAGGGGACCGTCCCCTGGGTAATGGCGAATGGG
ACC

### Protocol for the compound screening

#### Materials

- HEK 293T cells.
- T75 flask for adherent cells (TRP).
- Lipofectamine 2000 (Invitrogen).
- OptiMEM (Gibco).
- DMEM supplemented with 10% FCS.
- Trypsin-EDTA.
- Plasmids pcDNA6.B-nsp7-flag, pcDNA6.B-nsp8-flag, pcDNA6.B-nsp12-flag, pPolI-SARS-CoV2-NLuc-N.
- *Xba*I (NEB).
- PCR clean-up kit (NEB).
- RIPA lysis buffer (50mM Tris pH 7.8, 150mM NaCl, 1% NP-40, 0.5% sodium deoxycholate, 0.1% SDS).
- 0.1M DTT (Invitrogen).
- Complete protease inhibitor cocktail (Sigma).
- Nanoluciferase substrate (Promega).
- Centrifuge.
- 24-well plate (TPP).
- White 96-well plate (Greiner Bio-One).
- Luminometer (Promega).

#### Protocol

##### Cell suspension

- Split HEK 293T cells 1 day prior to experiment (note: using “old” cells, such as cells left to grow over the weekend, may increase variability and reduce reproducibility). Ideally, seed a T75 flask of HEK 293T cells, such that they will 80-90% on the day of the experiment. Incubate cells in DMEM/10% FCS at 37°C, 5% $CO_2$.
- On day of experiment, wash cells with 5ml PBS at RT.
- Trypsinize cells with 1 mL of tryspin-EDTA and incubate at 37°C until the cells detach from the flask.
- Inactivate trypsin with 4 mL of DMEM/10% FCS.
- Transfer the cell suspension to a 15 mL tube and centrifuge 5 min at 700 g.
- Resuspend cells in 5 mL of DMEM with 10% FCS.

##### Transfection and drug treatment

- Linearize 1 µg of pPolI-SARS-CoV2-NLuc-N with *Xba*I in rCutSmart buffer for 1 hour at 37°C.
- Clean-up digestion using PCR clean-up kit (NEB). Elute and adjust concentration to 25 ng/µl.
- For a transfection in a 24-well, prepare the following plasmid mix in a 1.5 mL tube:

    - 200 ng pcDNA6.B-nsp7-flag
    - 200 ng pcDNA6.B-nsp8-flag
    - 200 ng pcDNA6.B-nsp12-flag
    - 25 ng of linear pPolI-SARS-CoV2-NLuc-N
    - Add up to 10 µL OptiMEM.

- For a transfection in a 24-well, prepare the following lipofectamine mix in a 1.5 mL tube:

    - 108 µL of OptiMEM
    - 2 µL of lipofectamine 2000

- Incubate the lipofectamine/OptiMEM mix 5 min at room temperature.
- Add the lipofectamine mix to the plasmid mix and homogenize by pipetting.
- Centrifuge gently for a few seconds and incubate 20 min at room temperature.
- For a transfection in a 24-well, prepare the following cell mix directly in the plate:

    - 50 µL of HEK 293T cell suspension
    - 330 µL of DMEM/10% FCS.
    - 120 µl plasmid/lipofectamine

- Gently shake the plate and incubate at 37°C, 5% $CO_2$ for 3h.
- Three hours after transfection add antiviral compound directly in the cell supernatant.
- Gently shake the plate and incubate at 37°C, 5% $CO_2$ for 24h.

**Cell lysis and nanoluciferase assay**

- Place RIPA lysis buffer on ice and add fresh 1X protease inhibitor cocktail and 1 mM DTT.
- Remove the supernatant from the 24-well and add 200 µL cold complete RIPA lysis buffer.
- Place the 24-well plate on ice and incubate 5 min.
- Collect cell lysate in a 1.5 mL pre-chilled tube and homogenize by pipetting.
- Leave on ice for 10 min. Cells can be stored at -20°C at this stage.
- Spin down lysed cells and transfer 80 µL supernatant to a white 96-well plate.
- Add equal volume (80 µL) of nanoluciferase substrate.
- Shake plate and incubate lysed cells and substrate for 10 min at RT.
- Read luminescence using a luminometer with 0.5 sec integration time.

### Supplemental tables

##### Table S1. Primers for Nsp12 PCR mutagenesis

| nsp | Mutation | Forward primer | Reverse primer |
|---|---|---|---|
| nsp12 | DD761-762AA | TGATGATACTCTCTGGCAGCAGCTGTTGTGTGTTTC | TTTGTTCGAACCGCGGCTTATCGTCGTCATCCTTG |
| nsp12 | D619A | CACCTTATGGGTTGGGCTTATCCTAAATGTGATAG | CTATCACATTTAGGATAAGCCCAACCCATAAGGTG |
| nsp12 | P323L | ACAGTGTTCCCACTTACAAGTTTTGGA | TCCAAAACTTGTAAGTGGGAACACTGT |
| nsp12 | P322L | TCTCTACAGTGTTCCTACCTACAAGTTTTGG | CCAAAACTTGTAGGTAGGAACACTGTAGAGA |
| nsp7 | S25L | CTCAGAGTAGAATCATTATCTAAATTGTG | CACAATTTAGATAATGATTCTACTCTGAG |
| nsp7 | S26F | AGAGTAGAATCATCATTTAAATTGTGGG | CCCACAATTTAAATGATGATTCTACTCT |
| nsp8 | M129I | GCAGCCAAACTAATTGTTGTCATACCAGA | TCTGGTATGACAACAATTAGTTTGGCTGC |
| nsp8 | D163L | AGGTTGTAGATGCACTTAGTAAAATTGTTCA | TGAACAATTTTACTAAGTGCATCTACAACCT |

##### Table S2. PCR primers

| Gene name | Forward primer | Reverse primer |
|---|---|---|
| *Nsp7* | TTTTGCTTTCCATGCAGGGTG | TAAGGTTGCCCTGTTGTCCA |
| *Nsp8* | ACCTCTTACAACAGCAGCCAA | ATTTGACAGCAGAATTGGCCC |
| *Nsp9* | GTGCTGCCGGTACTACACAA | ACGTACTGTGGCAGCTAAACT |
| *Nsp10* | GTGGGGGACAACCAATCACT | ACGATGCACCACCAAAGGAT |
| *Nsp12* | AATAGAGCTCGCACCGTAGC | CCGCCACACATGACCATTTC |
| *Nsp13* | GCCGCTGTTGATGCACTATG | CTCCAAGCAGGGTTACGTGT |
| *Nsp14* | TGGGCACATGGCTTTGAGTT | GGAAAAGCATGTGGCACGTC |
| *Nsp16* | AAGACAGTGGTTGCCTACGG | TCGCGTGGTTTGCCAAGATA |
| *Nucleocapsid negative sense* | ACGAGAAGAGGCTTGACTGC | GTCTTGGTTCACCGCTCTCA |
| *Nucleocapsid positive sense* | TCTTGGTTCACCGCTCTCAC | TTGTTAGCAGGATTGCGGGT |
| *GAPDH* | CCACTAGGCGCTCACTGTTC | TGAGGTCAATGAAGGGGTCA |

**Table S3. Sequence of nsp12, nsp7 and nsp8 proteins**.

| **SARS-Cov-2 protein** | **Amino acid sequence** |
|---|---|
| nsp12<br><br>(932 aa) | SADAQSFLNRVCGVSAARLTPCGTGTSTDVVYRAFDIYNDKVAGFAKFLKTNCCRFQ<br>EKDEDDNLIDSYFVVKRHTFSNYQHEETIYNLLKDCPAVAKHDFFKFRIDGDMVPHIS<br>RQRLTKYTMADLVYALRHFDEGNCDTLKEILVTYNCCDDDYFNKKDWYDFVENPDIL<br>RVYANLGERVRQALLKTVQFCDAMRNAGIVGVLTLDNQDLNGNWYDFGDFIQTTP<br>GSGVPVVDSYYSLLMPILTLTRALTAESHVDTDLTKPYIKWDLLKYDFTEERLKLFDRYF<br>KYWDQTYHPNCVNCLDDRCILHCANFNVLFSTVF**PP**TSFGPLVRKIFVDGVPFVVST<br>GYHFRELGVVHNQDVNLHSSRLSFKELLVYAADPAMHAASGNLLLDKRTTCFSVAAL<br>TNNVAFQTVKPGNFNKDFYDFAVSKGFFKEGSSVELKHFFFAQDGNAAISDYDYYRY<br>NLPTMCDIRQLLFVVEVVDKYFDCYDGGCINANQVIVNNLDKSAGFPFNKWGKARL<br>YYDSMSYEDQDALFAYTKRNVIPTITQMNLKYAISAKNRARTVAGVSICSTMTNRQF<br>HQKLLKSIAATRGATVVIGTSKFYGGWHNMLKTVYSDVENPHLMGWDYPKCDRA<br>MPNMLRIMASLVLARKHTTCCSLSHRFYRLANECAQVLSEMVMCGGSLYVKPGGT<br>SSGDATTAYANSVFNICQAVTANVNALLSTDGNKIADKYVRNLQHRLYECLYRNRDV<br>DTDFVNEFYAYLRKHFSMMILSDDAVVCFNSTYASQGLVASIKNFKSVLYYQNNVF<br>MSEAKCWTETDLTKGPHEFCSQHTMLVKQGDDYVYLPYPDPSRILGAGCFVDDIVK<br>TDGTLMIERFVSLAIDAYPLTKHPNQEYADVFHLYLQYIRKLHDELTGHMLDMYSVM<br>LTNDNTSRYWEPEFYEAMYTPHTVLQ |
| nsp7<br><br>(83 aa) | SKMSDVKCTSVVLLSVLQQLRVES**SS**KLWAQCVQLHNDILLAKDTTEAFEKMVSLLS<br>VLLSMQGAVDINKLCEEMLDNRATLQ |
| nsp8<br><br>(198 aa) | AIASEFSSLPSYAAFATAQEAYEQAVANGDSEVVLKKLKKSLNVAKSEFDRDAAMQR<br>KLEKMADQAMTQMYKQARSEDKRAKVTSAMQTMLFTMLRKLDNDALNNIINNA<br>RDGCVPLNIIPLTTAAKL**M**VVIPDYNTYKNTCDGTTFTYASALWEIQQVVDA**D**SKIVQ<br>LSEISMDNSPNLAWPLIVTALRANSAVKLQ |

The amino acids shown in figure 4B-C are indicated in red.

### Supplemental figures

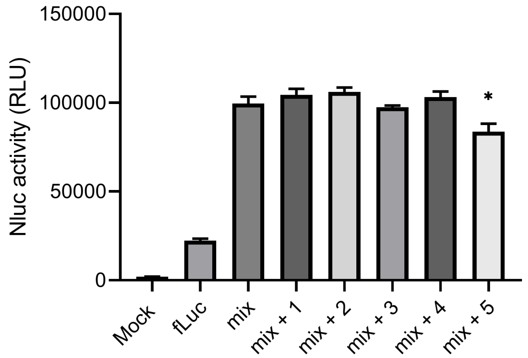


**Figure S1. Effect of additional of additional plasmid to Nluc signal.** Empty pcDNA3 plasmid was added to the nsp12, nsp7, nsp8 and Nluc transfection mix. Numbers indicate fold increases of the standard amount added. Bars represent mean of three biological repeats. Error bars indicate SEM. *P < 0.05; as determined by Dunnett's multiple comparisons test.

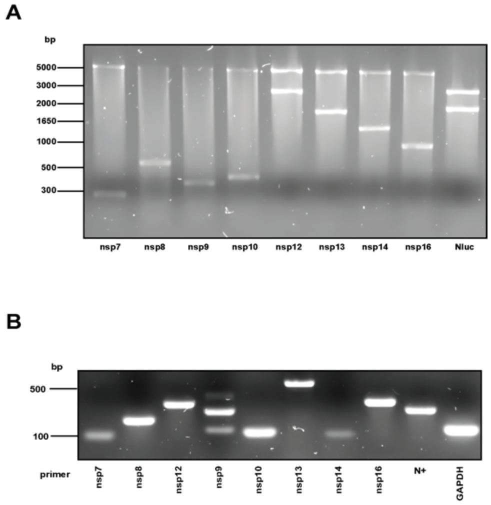


**Figure S2. Characterisation of different SARS-Cov-2 nsp plasmids.** (**A**) Double-digested plasmids containing SARS-CoV-2 nsp7, nps8, nsp9, nsp10, nsp12, nsp13, nsp14, nsp16 with BamHI/SacII and Nluc reporter plasmid with XbaI/StuI. (**B**) RT-PCR analysis of nsp7-flag, nsp8-flag, nsp12-flag, nsp9-flag, nsp10-flag, nsp13-flag, nsp14-flag, nsp16-flag, and positive sense Nluc-N gene expression 24 hours after transfection with nsp12-7-8-10-14-9-14-16 plasmids in HEK293T cells. GAPDH mRNA was detected as loading control. (**C**) Expression of positive-sense (+) reporter RNAs as determined by RT-PCR 24 hours after transfection. GAPDH mRNA was detected as loading control.
